# Kappa opioid receptors in excitatory spinal neurons gate acute pain: evidence from mouse and human

**DOI:** 10.64898/2026.07.01.735618

**Authors:** Nicole A. Kooij, Brandon M. Wills, Laney G. Waydick, Louis G. Fanien, Allison P. Manalo, Tayler D. Sheahan

**Affiliations:** Department of Cell Biology, Neurobiology, and Anatomy, Medical College of Wisconsin, Milwaukee, Wisconsin; Pittsburgh Center for Pain Research and Department of Neurobiology, University of Pittsburgh, Pittsburgh, Pennsylvania; Neuroscience Research Center, Medical College of Wisconsin, Milwaukee, Wisconsin

## Abstract

The kappa opioid receptor (KOR) has emerged as a promising, nonaddictive analgesic target, yet the neural mechanisms underlying KOR inhibition of pain are not entirely understood. Here, we provide converging evidence that KOR expressed on spinal neurons inhibits acute pain. We demonstrate that pharmacological inhibition of KOR-expressing neurons in the spinal cord blocks nocifensive behaviors. Conversely, chemogenetic activation of KOR-expressing spinal neurons elicits nocifensive behaviors. We then perform a series of molecular characterizations and show that excitatory KOR spinal neurons are recruited by noxious stimuli and coexpress pain-promoting neuropeptides such as Tac1 in both mouse and human. Together, these data suggest that kappa opioids inhibit pain by reducing the release of pain-promoting neuropeptides from a translationally relevant population of spinal neurons.

**In Brief:** Kooij et al. find KOR signaling within the spinal cord suppresses pain transmission. Molecular characterization of KOR-expressing spinal neurons reveals this may occur through the inhibition of excitatory spinal neurons, which also express pain-promoting neuropeptides.

**Highlights:**

- Spinal KOR signaling suppresses behavioral responses in rodent acute pain models
- Chemogenetic activation of *Oprk1^Cre^*spinal neurons elicits nocifensive behaviors
- KOR is expressed on excitatory superficial dorsal horn neurons that contain the pain-promoting neuropeptide Tac1 (Substance P) in mouse
- KOR-expressing spinal neurons similarly reside in the superficial dorsal horn and coexpress TAC1 in human

**Graphical Abstract:** 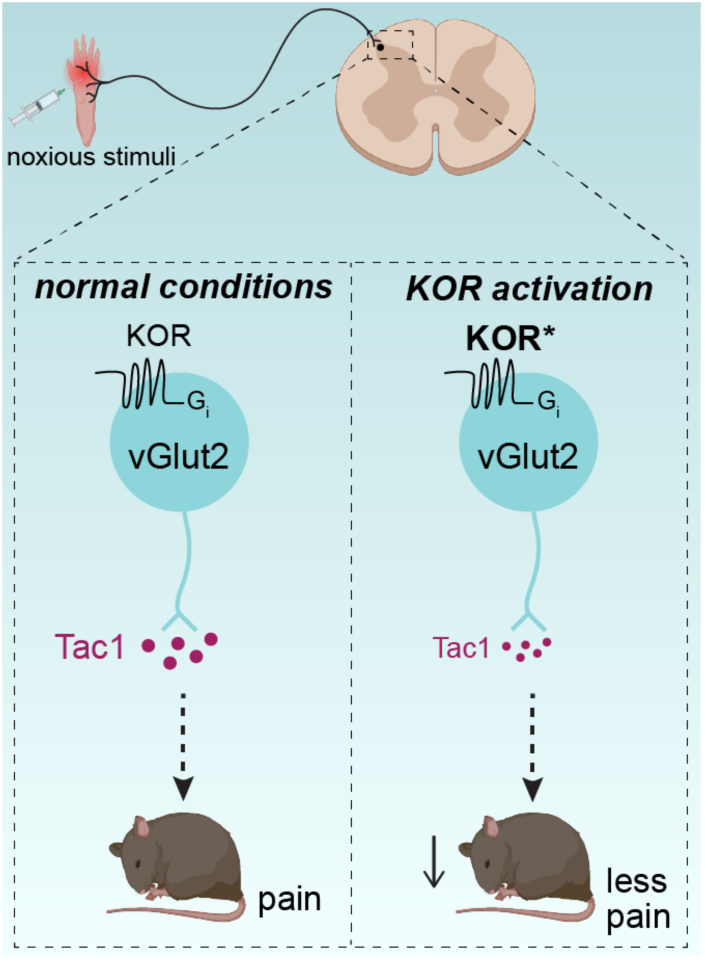

## Introduction

Opioids and their receptors are broadly expressed throughout the nervous system and profoundly shape nociception and pain^1–3^. To harness the analgesic potential of opioid receptors, synthetic opioids targeting the *mu* opioid receptor were developed and found to provide robust pain relief^4^. However, the addictive properties of *mu* opioids have contributed to the ongoing, worldwide opioid epidemic^5,6^. Alternative, nonaddictive analgesics are therefore highly sought after, which has led to an increasing interest in clinically targeting the kappa opioid receptor (KOR). Excitingly, KOR agonists show promise in the treatment of post-surgical, visceral, and abdominal pain^7,8^. However, our understanding of how the kappa opioid system shapes pain circuitry is incomplete.

KOR, an inhibitory G_i_-coupled GPCR, is expressed in both the peripheral and central nervous systems and is therefore well-positioned to influence pain at multiple levels. Interestingly, KOR has been shown to differentially modulate pain depending on where within the nervous system it is activated. In the periphery, previous work has established that KOR is expressed in nociceptive sensory neurons, and KOR activation reduces inflammatory and visceral pain^9–13^. In contrast, within the brain, KOR signaling acts on descending circuits to facilitate pain^14–17^. Moreover, recruitment of the kappa opioid system within limbic circuits has been shown to promote the negative affect that accompanies pain states^18–22^. While KOR inhibition of pain has been studied extensively in the peripheral nervous system and supraspinal circuits, we lack detailed mechanistic insight into how KOR influences pain within the spinal cord.

We and others have shown that both KOR and its endogenous ligand, dynorphin, are expressed within the spinal cord dorsal horn^23–27^. However, there is conflicting evidence as to whether activation of the kappa opioid receptor is analgesic within the spinal cord. Early studies in which dynorphin was delivered locally to the spinal cord via intrathecal injection demonstrated that dynorphin elicited allodynia, suggesting the spinal kappa opioid system facilitates pain^28,29^. In contrast, intrathecal delivery of selective KOR agonists has been shown to be antinociceptive^30,31^. These data have been difficult to reconcile because the spinal neuron populations that express KOR, and in turn where kappa opioids act within spinal cord dorsal horn circuits, have not been characterized.

Here, we address these major unresolved questions surrounding how the kappa opioid system influences pain at the spinal level. We demonstrate that inhibition of KOR-expressing neurons within the spinal cord attenuates acute pain. Conversely, activation of KOR-expressing spinal neurons is sufficient to elicit pain-related behaviors. To provide mechanistic insight into how KOR signaling causes these effects, we performed the first in-depth molecular characterization of KOR-expressing spinal neurons and show that excitatory KOR spinal neurons convey painful stimuli, and that these neurons also express pain-promoting neuropeptides including *Tac 1* (Substance P). Lastly, we show that a similar population of neurons exists within the human spinal cord, highlighting the translational relevance of our rodent findings. Together, these studies provide a detailed picture of KOR expression and function within the spinal cord dorsal horn.

## Results

### The kappa opioid receptor inhibits acute pain within the spinal cord

We first asked whether KOR signaling modulates pain within the spinal cord using pharmacological strategies (Fig. 1A). As KOR signaling decreases neuronal activity (Fig. 1B), we hypothesized that activation of KOR on spinal neurons would decrease pain. To examine this, we used the clinically effective KOR agonist nalfurafine, which we recently showed suppresses itch by acting on spinal neurons^25^. In keeping with these results, intrathecal injection of nalfurafine reduced nocifensive behaviors in response to intraplantar capsaicin and formalin (Fig. 1C, D), demonstrating that spinal KOR signaling blocks pain behaviors.

**Figure 1.**
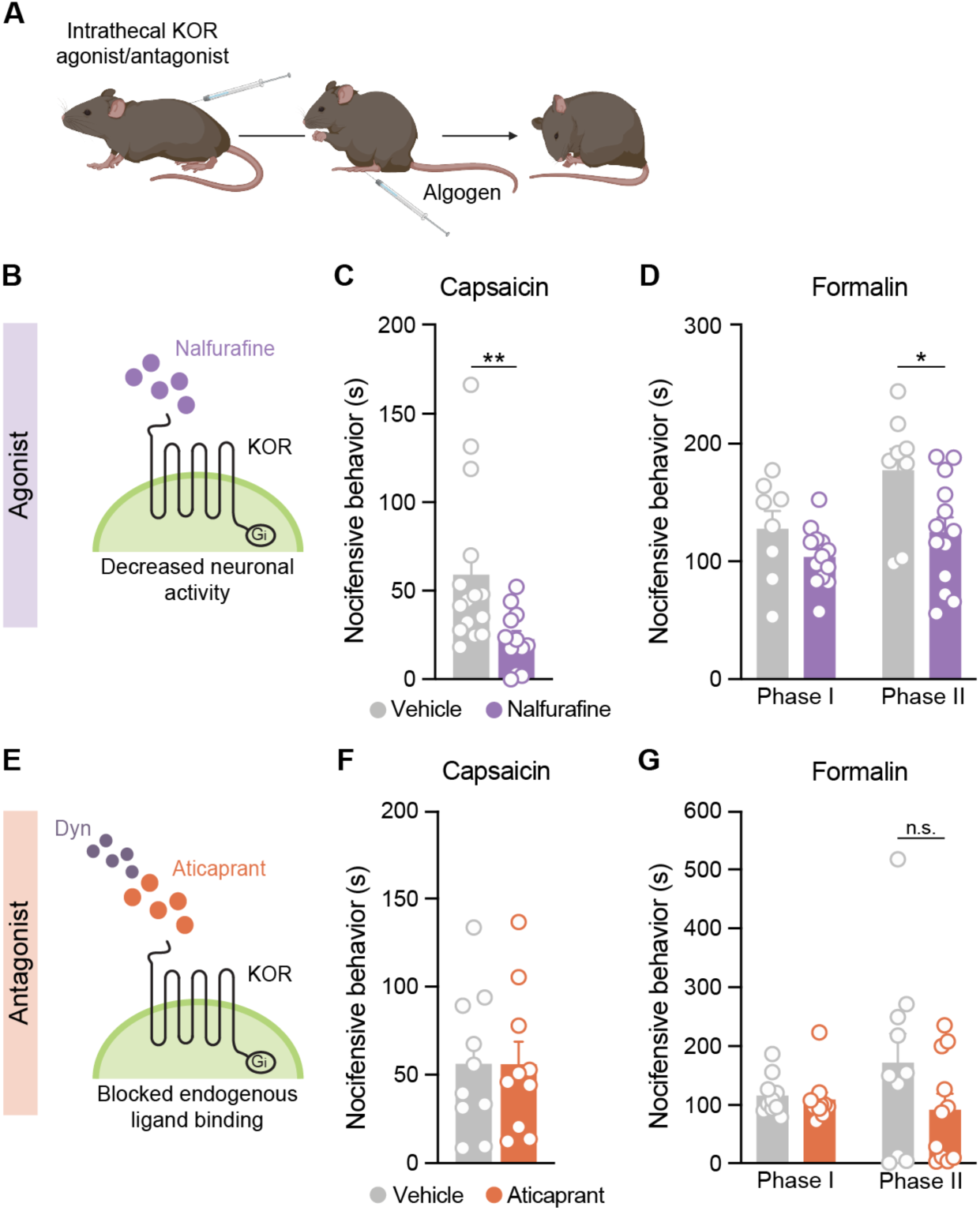
Pharmacological activation of the kappa opioid receptor within the spinal cord suppresses acute pain behavior. **(A)** Mice were first injected intrathecally with either a KOR agonist, antagonist, or vehicle, followed by an intraplantar algogen injection 30 min later, and the duration of nocifensive behaviors was quantified. **(B)** Schematic showing the mode of action of the KOR agonist nalfurafine on KOR-expressing neurons, resulting in decreased neuronal activity. **(C)** Intrathecal administration of nalfurafine (40 µg) significantly reduced capsaicin-induced nocifensive behavior (n = 12-15 mice per group; Mann-Whitney test) and **(D)** Phase II of formalin-induced nocifensive behavior (n = 8-13 mice per group; 2-way RM ANOVA, main effect of nalfurafine: *p<0.05) relative to pretreatment with vehicle. **(E)** Schematic showing the mode of action of the KOR antagonist aticaprant (10 µg) on KOR-expressing neurons, resulting in blockade of the binding of endogenous ligands, such as dynorphin. **(F)** Intrathecal administration of aticaprant did not affect capsaicin-induced nocifensive behavior (n = 10 mice per group; Student’s unpaired t-test) or **(G)** formalin-induced nocifensive behavior (n = 10-11 mice per group; 2-way RM ANOVA, no main effect of aticaprant) as compared to vehicle-treated mice. Data are shown as mean ± SEM, with open circles representing individual mice. *p<0.05, **p<0.01, n.s. not significant.

The spinal circuits that convey pain are closely related to those that convey itch. In previous studies examining the role of the kappa opioid system in the spinal coding of itch, intrathecal kappa antagonists elevated scratching responses to pruritogens, highlighting that the endogenous KOR ligand dynorphin gates spinal transmission of itch^46,47^. Thus, we next asked whether endogenous KOR signaling also gates pain behaviors (Fig. 1E). Intrathecal injection of the KOR antagonist aticaprant did not alter nocifensive responses to intraplantar capsaicin or formalin (Fig. 1F, G), suggesting that dynorphin may differentially modulate acute pain and itch. Together, these results demonstrate that KOR signaling inhibits acute pain within the spinal cord.

### *Oprk1* is expressed throughout the spinal cord dorsal horn of mouse and human

After demonstrating that activation of KOR within the spinal cord suppresses acute pain, our overarching goal was to understand which spinal neurons KOR agonists act on to reduce pain. As a first step, we asked where KOR is expressed within the mouse spinal cord. We performed fluorescence *in situ* hybridization (FISH) on wild-type mice and probed for *Oprk1*, the gene encoding KOR. RNA-sequencing studies indicate that kappa opioid receptor expression is largely absent from non-neuronal cells in the spinal cord across species, and thus positive cells were reasonably assumed to be neurons^48–50^. We observed a broad laminar distribution of *Oprk1* neurons across the mouse dorsal horn, including the superficial dorsal horn (37% of *Oprk1*^+^ neurons), deep dorsal horn (59% of *Oprk1*^+^ neurons), and a small subset of neurons in the lateral spinal nucleus (4% of *Oprk1*^+^ neurons) (Fig. 2A-C).

**Figure 2.**
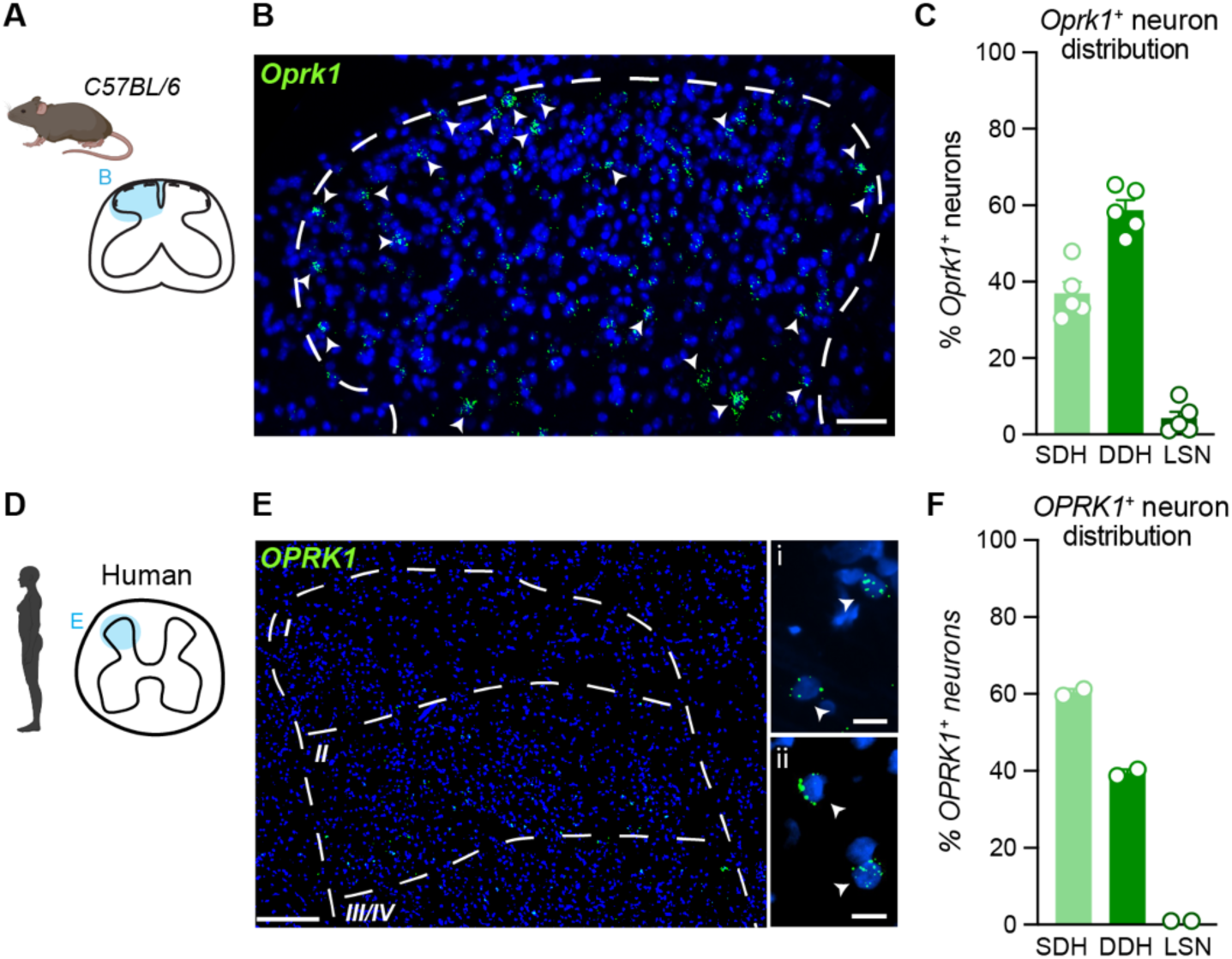
Distribution of *Oprk1*-expressing neurons in the mouse and human spinal cord dorsal horn. **(A)** Expression of *Oprk1* was visualized in the spinal cord dorsal horn of C57BL/6 mice using FISH. **(B)** Representative image of *OPRK1*-expressing neurons in the mouse dorsal horn. Scale bar, 50 µm. **(C)** Quantification of the percentage of *Oprk1^+^* neurons distributed throughout the mouse superficial dorsal horn (SDH), deep dorsal horn (DDH), and lateral spinal nucleus (LSN). N=5 mice; n = 67-133 total *Oprk1* neurons per mouse. **(D)** Expression of *OPRK1* was visualized in the human spinal cord dorsal horn using FISH. **(E)** Representative image of *OPRK1*-expressing neurons in the human superficial dorsal horn. Scale bars: E, 250 μm; E_i_ and E_ii_, 25 μm. **(F)** Quantification of the percentage of *OPRK1^+^* neurons distributed throughout the human dorsal horn. N = 2 human donors; n = 76-99 total *OPRK1* neurons per donor. B, E: Arrowheads, *Oprk1*^+^ neurons. Data are shown as mean ± SEM, with open circles representing individual mice or humans.

To assess the translational relevance of our rodent findings, we next asked whether KOR might also be well-positioned within the human spinal cord to modulate pain. We conducted analogous FISH analyses and examined the distribution of *OPRK1* neurons in spinal cord samples from human organ donors. As in mouse, *OPRK1* neurons were present throughout the human dorsal horn, including the superficial dorsal horn (60.5% of *OPRK1*^+^ neurons) and deep dorsal horn (39.5% of *OPRK1*^+^ neurons). No *OPRK1*^+^ neurons were detected in the lateral spinal nucleus of human (Fig. 2D-F, S1A). These results show that KOR is present in the superficial dorsal horn of both mouse and human and is therefore well-positioned to suppress pain in both species.

### Activation of *Oprk1^Cre^*spinal neurons increases acute pain

Our findings that KOR agonists suppress pain when delivered intrathecally and that *Oprk1* is expressed on neurons within the superficial dorsal horn suggested KOR spinal neurons may encode pain. However, KOR is also known to be expressed on nociceptive sensory neurons that innervate the superficial dorsal horn^9,51,52^. Thus, the finding that intrathecally administered KOR agonists suppress acute pain could reflect inhibition of either sensory neuron central terminals or spinal neurons. To determine whether KOR spinal neurons drive pain, we selectively targeted KOR-expressing spinal neurons using an *Oprk1^Cre^* allele coupled with intraspinal viral injections. First, we confirmed that *Oprk1^Cre^*expression aligned with endogenous expression of *Oprk1* by injecting a Cre-dependent virus encoding nuclear-restricted EGFP, enabling us to visualize *Oprk1^Cre^* spinal neurons (Oprk1::nucEGFP, AAV8-EF1a-Nuc-flox(mCherry)-EGFP). As with *Oprk1* expression in wild-type mice, the majority of virally-labeled *Oprk1^Cre^* spinal neurons resided within the superficial dorsal horn (55.8% of *Oprk1^Cre^* neurons) and deep dorsal horn (43.3% of *Oprk1^Cre^* neurons) (Fig. S2A-S2C), highlighting the *Oprk1^Cre^* allele as a powerful tool to study spinal neuron function.

We then asked whether chemogenetic activation of *Oprk1^Cre^*spinal neurons could elicit pain-related behaviors. To selectively activate KOR spinal neurons, we injected Cre-dependent viruses encoding either an excitatory DREADD fused to a fluorescent reporter (Oprk1::hM3Dq, AAV8-hSyn-DIO-hM3Dq-mCherry) or a Cre-dependent control virus containing only a fluorophore (Oprk1::mCherry, AAV8-hSyn-DIO-mCherry) into the lumbar spinal cord of *Oprk1^Cre^* mice (Fig. 3A). These injections provided robust labeling of *Oprk1^Cre^* spinal neurons and their processes throughout the dorsal horn (Fig. 3B, S3A). No viral labeling was detected in lumbar dorsal root ganglia neurons (Fig. 3C), ensuring that our chemogenetic manipulations were specific to *Oprk1^Cr^*^e^ spinal neurons.

**Figure 3.**
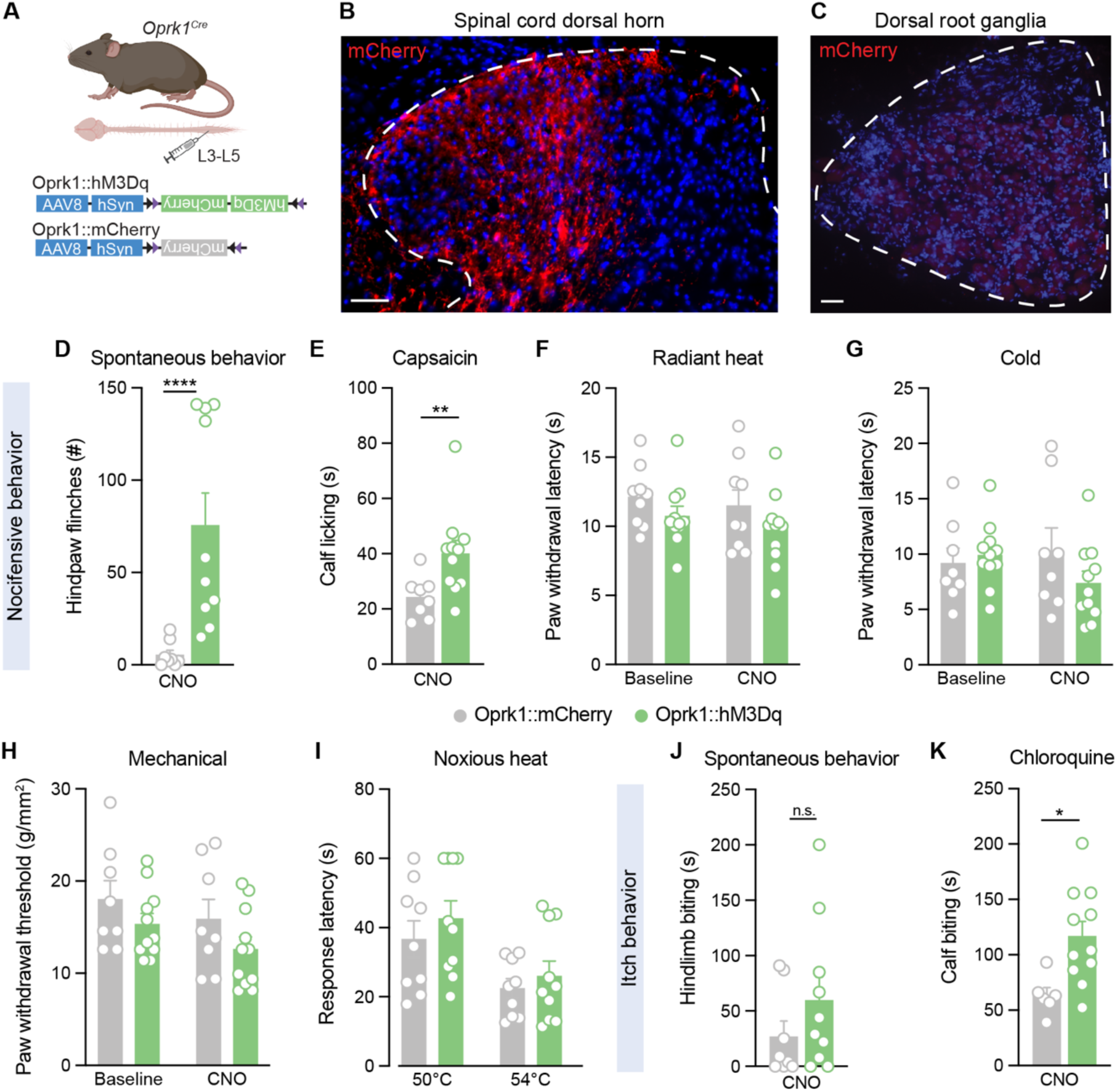
Chemogenetic activation of *Oprk1* spinal neurons increases nocifensive behaviors. **(A)** Strategy to selectively target *Oprk1^Cre^*spinal neurons. To gain chemogenetic access to *Oprk1* spinal neurons, the Cre-dependent viruses AAV8-hSyn-DIO-hM3Dq-mCherry (Oprk1::hM3Dq) or AAV8-hSyn-DIO-mCherry (Oprk1::mCherry) were injected unilaterally into the lumbar spinal cord of *Oprk1^Cre^* mice. **(B)** Virally-mediated recombination, as indicated by mCherry expression, was present in the spinal cord dorsal horn, **(C)** but not the corresponding lumbar dorsal root ganglia of *Oprk1^Cre^* mice. Images representative of n = 3 mice. Scale bars, 50 µm. **(D)** Chemogenetic activation of *Oprk1^Cre^* spinal neurons (CNO, 5 mg/kg, i.p.) elicited spontaneous hindpaw flinches, a nocifensive behavior, in Oprk1::hM3Dq mice, but not Oprk1::mCherry controls (n = 8-10 mice per group). Mann-Whitney test. **(E)** Activation of *Oprk1^Cre^*spinal neurons increased the duration of site-directed licking in response to intradermal injection of capsaicin (0.001%, 10 µL) into the calf (n = 8-11 mice per group). Mann-Whitney test. **(F)** Hindpaw nociceptive withdrawal thresholds to radiant heat (n = 9-11 mice per group; 2-way RM ANOVA) **(G)** cold (n = 8-11 mice per group; 2-way RM ANOVA), and **(H)** mechanical stimuli (n = 8-11 mice per group; 2-way RM ANOVA) were unchanged in Oprk1::hM3Dq mice following CNO administration. **(I)** Response latencies to noxious heat did not change in response to activation of *Oprk1^Cre^*spinal neurons (n = 9-10 mice per group). Unpaired Student’s T-test, Holm-Sidak correction for multiple comparisons. **(J)** Chemogenetic activation of *Oprk1^Cre^* spinal neurons alone did not alter site-directed biting of the hindlimb, an itch-related behavior (n = 8-10 mice per group). Unpaired Student’s T-test. **(K)** Activation of *Oprk1^Cre^*spinal neurons increased the duration of site-directed biting in response to intradermal injection of chloroquine (100 µg/5 µL) into the calf. (n = 6-11 mice per group). Unpaired Student’s T-test. Data are shown as mean ± SEM, with open circles representing individual mice. n.s. not significant, *p<0.05, **p<0.01, ****p<0.0001

We then chemogenetically activated *Oprk1^Cr^*^e^ spinal neurons in naive mice and assessed changes in pain-like behaviors. Activation of *Oprk1^Cr^*^e^ spinal neurons caused spontaneous hindpaw flinching (Fig. 3D, S3B), a nocifensive behavior. Similarly, activation of *Oprk1^Cr^*^e^ spinal neurons increased nocifensive responses to intraplantar capsaicin (Fig. 3E). In contrast, in stimulus-evoked somatosensory assays, chemogenetic activation of *Oprk1^Cr^*^e^ spinal neurons did not alter nociceptive responses to heat, cold, or mechanical stimuli (Fig. 3F-I). Overall, these findings demonstrate that KOR neurons transmit noxious signals within the spinal cord.

In some studies, DREADDs have been reported to have constitutive activity, and in turn alter baseline nociceptive thresholds^53,54^. To assess this possibility, we also measured the baseline thresholds of Oprk1::mCherry and Oprk1::hM3Dq mice in the absence of CNO. Expression of hM3Dq alone did not affect baseline sensitivity to heat, cold, or mechanical stimuli compared with expression of the control virus (Fig. 3F-H). CNO administration alone similarly had no effect on nociceptive withdrawal thresholds, as responses thresholds of the hindpaw contralateral to intraspinal viral injection in Oprk1::mCherry or Oprk1::hM3Dq mice were unchanged following CNO relative to baseline (Fig. S3C -S3E).

### Activation of *Oprk1^Cr^*^e^ spinal neurons increases acute itch

Through primarily pharmacological approaches, we and others have demonstrated that KOR signaling blocks itch within the spinal cord^25,27,46,47,55^. To confirm and extend these findings, we next asked whether chemogenetic activation of *Oprk1^Cr^*^e^ spinal neurons drives itch behaviors. Surprisingly, we found no significant change in spontaneous itch behavior (Fig. 3J). One possible explanation for this is that animals were displaying spontaneous nocifensive flinches at this time (Fig. 3D), which could support the idea of salience processing of ascending somatosensory inputs^56^. However, chemogenetic activation of *Oprk1^Cr^*^e^ spinal neurons potentiated behavioral responses to intradermal injection of the pruritogen chloroquine (Fig. 3K). These findings are consistent with our previous work^25^ and underscore the importance of KOR neurons in the spinal transmission of itch.

### Excitatory *Oprk1* spinal neurons convey noxious stimuli

Having found that activating KOR spinal neurons drives pain behavior, our next goal was to characterize which pain-encoding neurons in the spinal cord express KOR. As a first step towards answering this question, we asked whether *Oprk1* is expressed in excitatory or inhibitory spinal neurons by performing FISH on wild-type mice (this data set was also analyzed in a previous publication^25^, and we extend these analyses here). We found that *Oprk1* spinal neurons are composed of both excitatory (*Vglut2^+^,* 43%) and inhibitory (*Vgat^+^,* 49%) neurons (Fig. 4A, B), consistent with recent RNA-sequencing data sets of spinal neurons^49^. Based on this finding, we reasoned that there are two models by which activating KOR spinal neurons could elicit pain behavior (Fig. 4C). In one model (facilitation), KOR-expressing excitatory spinal neurons drive pain. In the other model (disinhibition), KOR-expressing inhibitory spinal neurons ungate pain by disinhibiting the excitatory neurons that drive pain.

**Figure 4.**
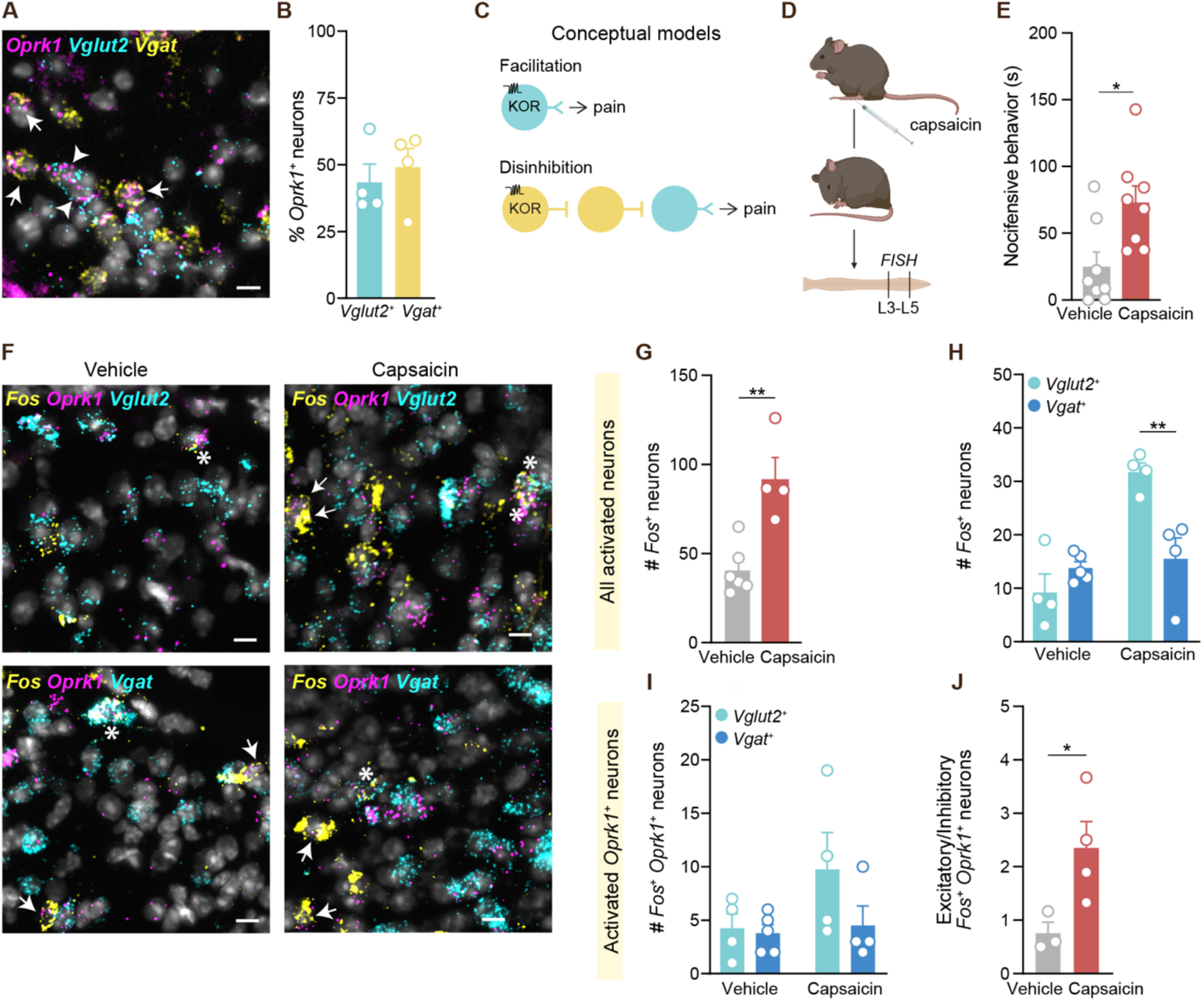
Excitatory *Oprk1* spinal neurons convey acute pain. **(A)** Representative FISH image demonstrating expression of *Vglut2* and *Vgat* within *Oprk1* neurons in the superficial dorsal horn. Scale bar, 10 µm. **(B)** Quantification of the percentage of *Oprk1^+^* SDH neurons that express *Vglut2* or *Vgat* (N=4 mice; n=45-71 total *Oprk1* neurons per mouse). **(C)** Conceptual models for how activation of KOR spinal neurons may cause pain-related behaviors. Activation of excitatory KOR spinal neurons may facilitate pain (top), or activation of inhibitory KOR spinal neurons may disinhibit neurons that gate pain (bottom). **(D)** Mice were given an intraplantar injection of capsaicin (0.01%, 10 µL) or vehicle, and the duration of nocifensive behaviors was quantified for 15 min. The lumbar spinal cord from approximately L3-L5 was dissected and used for FISH analysis. **(E)** Intraplantar injection of capsaicin significantly increased nocifensive behavior compared to vehicle (n = 8 mice per group). Unpaired Student’s T-test. **(F)** Representative FISH images of superficial dorsal horn neurons activated in response to intraplantar injection of vehicle or capsaicin, as revealed by *Fos* expression, and the coexpression of *Vglut2*, *Vgat*, and/or *Oprk1* within these neurons. Scale bars, 10 µm. Arrow, *Fos*^+^ *Oprk1* neuron; *triple-labeled neuron. **(G)** The total number of *Fos*^+^ in the superficial dorsal horn was increased by intraplantar injection of capsaicin (n= 4-6 mice per group). Unpaired Student’s T-test. **(H)** Intraplantar capsaicin selectively activated excitatory (*Vglut2*^+^) neurons within the superficial dorsal horn (n= 4-5 mice per group). 2-way ANOVA, treatment x cell-type interaction: **p<0.01; main effect of cell-type: *p<0.05. Holm-Šídák’s correction for multiple comparisons. **(I)** The number of excitatory vs inhibitory *Fos*^+^ *Oprk1* neurons in the dorsal horn of mice treated with capsaicin versus vehicle (n= 4-5 mice per group). 2-way ANOVA, treatment x cell-type interaction: n.s. **(J)** The ratio of activated excitatory *Oprk1*^+^ neurons to activated inhibitory *Oprk1*^+^ neurons increased in response to capsaicin as compared to vehicle (n = 3-4 mice per group). Unpaired Student’s T-test. Data are shown as means ± SEM, open circles representing individual mice n.s. not significant, *p<0.05, **p<0.01.

To distinguish between these models, we visualized the neurons in the superficial dorsal horn that were activated by noxious stimuli. Following intraplantar injection of capsaicin or vehicle, we recorded nocifensive behaviors, then performed FISH and identified activated spinal neurons based on the expression of the immediate early gene *Fos* (Fig. 4D). As expected, intraplantar capsaicin elicited more nocifensive behaviors as compared to vehicle (Fig. 4E), which also coincided with an increase in the number of *Fos*^+^ spinal neurons within the superficial dorsal horn (Fig. 4F,G). We then asked whether excitatory or inhibitory spinal neurons were recruited by noxious input. Compared to vehicle conditions in which equivalent numbers of *Vglut2^+^*and *Vgat^+^* neurons expressed *Fos*, capsaicin caused a selective increase in the number of *Vglut2^+^ Fos* neurons (Fig. 4F, H). A similar trend was observed amongst *Oprk1*^+^ *Fos* neurons, such that capsaicin injection caused activation of more than two times as many excitatory *Oprk1* neurons than inhibitory *Oprk1* neurons as compared to vehicle (Fig. 4F, I, J). Based on these findings, the simplest model for how KOR spinal neurons drive pain is that increased activity of KOR-expressing excitatory neurons facilitates pain transmission within the spinal cord.

### *Oprk1* is expressed in excitatory dorsal horn neurons that contain pain-promoting neuropeptides in mouse and human

Lastly, we sought to further define the molecular identity of the excitatory *Oprk1* spinal neurons that encode pain. Populations of excitatory superficial dorsal horn neurons can be defined based on their expression of neuropeptide markers: *Tac1*, *Npff*, *Grp*, *Tac2*, *Cck*, and *Nts*^57^ (Fig. 5A). Thus, we used these neuropeptides as a framework to understand which excitatory neurons express *Oprk1*. We performed FISH on wild-type mice and quantified the percentage of excitatory *Oprk1* neurons in the superficial dorsal horn that expressed each neuropeptide marker (a portion of these dataset were also analyzed in a previous publication^25^, and we extend our analyses here). We found that excitatory *Oprk1* neurons express a variety of neuropeptides: *Tac1* (42%), *Npff* (33%), *Grp* (25%), *Tac2* (17%), and *Cck* (6%) (Fig. 5B-G). *Nts* was not detected within excitatory *Oprk1* neurons (Fig. 5B, H). When totaled, the percentages of *Oprk1* neurons that express each neuropeptide are more than 100% because while the neuropeptides mark largely distinct populations, there is some overlap in expression^57^. As neuropeptides are key signaling molecules in spinal pain circuits, this molecular profiling strategy also provided insights into how KOR signaling could shape pain spinal circuitry. Notably, several of the neuropeptides we found to be expressed in excitatory *Oprk1* spinal neurons are known to promote pain within the dorsal horn, including *Tac1*, *Tac2*, and *Cck*^58–60^.

**Figure 5.**
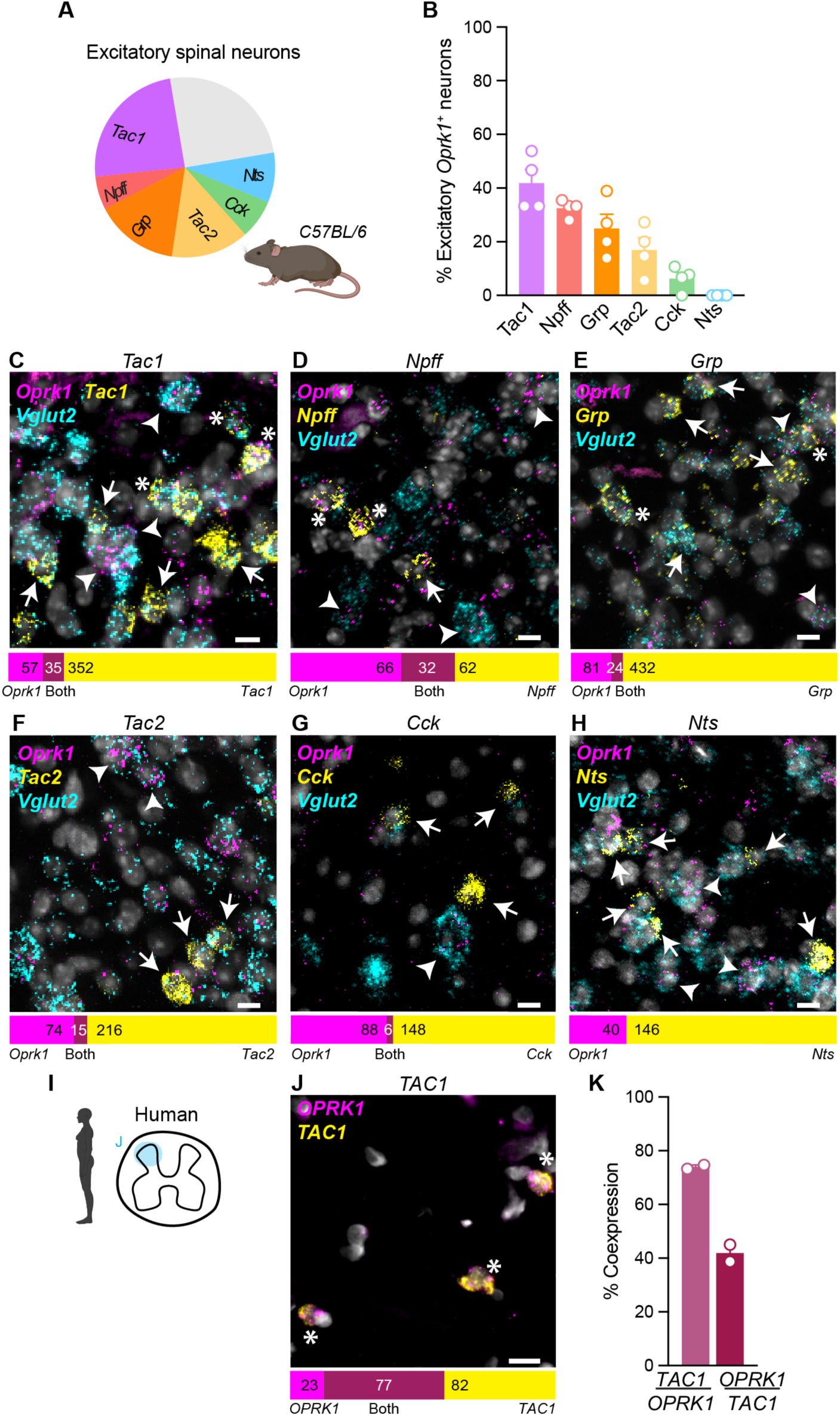
*Oprk1* is expressed in superficial dorsal horn neurons that contain pain-promoting neuropeptides in both mouse and human. **(A)** Pie chart showing the proportion of excitatory superficial dorsal horn spinal neurons that express established excitatory neuropeptide markers, adapted from Piers and Todd 2020^57^. **(B)** Quantification of the percentage of excitatory (*Vglut2*^+^) *Oprk1* superficial dorsal horn neurons that coexpress each neuropeptide in C57BL/6 mice (N = 4 mice per neuropeptide marker; *Oprk1* neurons per mouse: *Tac1*: n = 13-36, *Npff*: n = 15-31, *Grp*: n = 18-36, *Tac2*: n = 15-34, *Cck*: n = 13-30, *Nts*: n = 7-12). Representative images of excitatory *Oprk1* superficial dorsal horn neurons that express (**C**) *Tac1*, (**D**) *Npff*, (**E**) *Grp*, (**F**) *Tac2*, (**G**) *Cck*, or (**H**) *Nts*. Scale bars, 10 μm. Arrowhead, excitatory *Oprk1* neuron; arrow: neuropeptide-positive neuron; *triple-labeled neuron. **(I)** Expression of *OPRK1* was visualized in the human lumbar spinal cord dorsal horn using FISH. **(J)** Representative FISH image demonstrating coexpression of *OPRK1* and *TAC1* and within human superficial dorsal horn neurons. Scale bar, 50μm. *dual-labeled neurons. **(K)** Quantification of the coexpression of *OPRK1* and *TAC1* within superficial dorsal horn neurons (N= 2 donors; neurons per donor: *OPRK1*: n =59-46, *TAC1*: n =94-65). Data are shown as mean ± SEM, with open circles representing individual mice or humans.

We then asked whether kappa opioid signaling inhibits pain in the human spinal cord through a similar mechanism. For this to be the case, we would expect KOR to be expressed with the same pain-promoting neuropeptides in human. To examine this, we performed FISH on human spinal cord sections and asked whether *OPRK1* spinal neurons coexpress *TAC1*, the neuropeptide that was most prevalent in *Oprk1^+^* mouse spinal neurons (Fig. 5I). Excitingly, we found that the majority (74%) of *OPRK1* human superficial dorsal horn neurons coexpress *TAC1* (Fig. 5J-K). Conversely, 42% of *TAC1* neurons were *OPRK1*^+^ (Fig. 5J-K). Overall, these findings suggest that kappa agonists may block pain by reducing the activity of excitatory neurons that release pain-promoting neuropeptides in the spinal cord of mouse and human.

## Discussion

In this study, we revealed how the kappa opioid receptor system influences pain within the spinal cord. Using complementary behavioral pharmacology and chemogenetic approaches, we showed that, as in the periphery, kappa opioid receptor signaling reduces pain transmission at the spinal level. Further, we established the molecular identity of KOR neurons to provide mechanistic insight as to how spinal KOR signaling blocks pain inputs. We showed that KOR is expressed in excitatory spinal neurons that drive pain behaviors and contain neuropeptides that are known to propagate pain. Excitingly, KOR-expressing neurons in humans also contain these pain-promoting neuropeptides, raising the possibility that the mechanisms we identified here in mice are conserved in humans and therefore have translational relevance. Thus, our study provides comprehensive, cross-species insights into the functional and molecular identity of neurons in the spinal cord that express the kappa opioid receptor. More broadly, by building on prior efforts to map kappa opioid modulation of pain in both the periphery and the brain, this work provides the missing pieces of the puzzle needed to achieve a more complete understanding of KOR’s modulation of the pain across the neuraxis.

### KOR is found in excitatory spinal neurons that encode pain

Through a series of chemogenetic, behavioral, and molecular profiling approaches, our data indicate that while KOR is expressed in both excitatory and inhibitory spinal neurons, the excitatory subset of KOR neurons is recruited in response to noxious stimuli. Thus, the most likely mechanism by which KOR signaling suppresses acute pain is by dampening the activity of excitatory spinal neurons that encode pain. Nevertheless, what role inhibitory KOR spinal neurons play in somatosensation remains an outstanding question. Amongst other receptors known to be expressed in both excitatory and inhibitory spinal neuron populations, there is precedent for these cell-types working in parallel to facilitate changes in somatosensation. For instance, excitatory spinal neurons that express the *mu* opioid receptor have been proposed to mediate exogenous opioid-induced analgesia, while their inhibitory counterparts mediate endogenous opioid-induced analgesia^27,61,62^. Oxytocin-receptor expressing spinal neurons are similarly composed of half excitatory and inhibitory neurons, and these populations work in tandem to enhance the perception of pleasant touch^63^. Whether this framework extends to KOR inhibition of pain is an interesting question that merits further investigation.

Upon finding that KOR is expressed in excitatory spinal neurons, we categorized these cells based on their expression of neuropeptides. However, another way to understand how they fit into spinal pain circuits is by classifying them as interneurons or spinal output neurons. We recently showed that *Oprk1* is expressed on spinal output neurons that convey ascending somatosensory inputs to the brain. We found that ∼50% of all spinal output neurons within the superficial dorsal horn express *Oprk1*, and these target a number of structures important for pain processing including the parabrachial nucleus, periaqueductal gray, and caudal ventral posterolateral thalamus^25^. These observations raise the possibility that KOR agonists are effective in reducing acute pain in part because they act on spinal output neurons. This idea is supported by our finding that the increase in the number of activated excitatory *Oprk1* neurons under acute pain conditions (intraplantar capsaicin) is modest, yet pharmacological and chemogenetic manipulations of KOR and *Oprk1^Cre^* spinal neurons, respectively, have robust effects on behavioral responses to capsaicin. Future studies in which *Oprk1*^+^ spinal output neurons are selectively targeted are needed to determine whether they are a cellular substrate for spinal kappa opioid inhibition of pain.

### KOR signaling inhibits release of pain-promoting neuropeptides

Our results point to a mechanism in which KOR signaling within excitatory spinal neurons dampens pain transmission by reducing the release of key molecular mediators of pain, particularly neuropeptides such as Substance P (encoded by *Tac1*). Intriguingly, there is recurring evidence for kappa opioid disruption of neuropeptide release at multiple nodes within pain neural circuitry. In the periphery, KOR signaling inhibits neurogenic inflammation, a process mediated by the release of the neuropeptides Substance P and CGRP from sensory neuron peripheral terminals^9,64^. Moreover, previous studies in the spinal cord showed that activation of KOR similarly reduces local release of Substance P^65–67^. While these studies posited that kappa opioid signaling blocked neuropeptide release from primary afferents, our findings that *Oprk1* and *Tac1* are coexpressed in spinal neurons reveals that KOR is positioned to inhibit neuropeptide release from spinal neurons as well. Whether KOR signaling has similar effects in supraspinal brain structures that receive and integrate nociceptive inputs from the spinal cord is not yet known. In support of this possibility, existing evidence shows *Oprk1* is expressed within the parabrachial nucleus^68^, and KOR signaling appears to alter PBN neuron excitability^69^. Thus, the reduction of neuropeptide release is a recurrent mechanism by which kappa opioid receptor signaling shapes neural circuit function to reduce pain.

### KOR inhibition of pain in acute and chronic states

Previous investigations of whether KOR signaling in the spinal cord is analgesic have been controversial. Here, we settle this debate by demonstrating that selective KOR agonists reduce nocifensive behaviors at the spinal level in two models of acute pain: capsaicin and formalin. When considering early evidence that intrathecal dynorphin causes allodynia, our findings are consistent with the previously proposed idea that allodynic effects are mediated by actions at non-opioid receptors, such as NMDA^28,29^.

Having established that KOR signaling in spinal neurons blocks acute pain, our results beg the question of whether KOR agonists might also be effective in models of chronic pain. Several studies have demonstrated that intrathecal administration of KOR agonists reverses nerve injury-induced mechanical and thermal hypersensitivity^70–72^. Conversely, intrathecal delivery of KOR antagonists is sufficient to reinstate hypersensitivity after injuries have resolved^73,74^. These findings underscore that not only do exogenous kappa agonists block chronic pain, but that the endogenous dynorphin-kappa opioid receptor system is recruited to suppress chronic pain. In close alignment with this, upregulation of dynorphin’s precursor protein, prodynorphin, within the dorsal root ganglia and spinal cord is a proposed indicator of persistent pain states^75–77^.

A common issue with pharmacological management of chronic pain in patients is the development of analgesic tolerance. Therefore, to have clinical potential, it is critical to understand whether KOR agonists would also trigger the development of tolerance when used chronically. Biased KOR agonists have been shown to retain clinical efficacy with repeated dosing in chronic itch patients^78,79^, and evidence from preclinical models investigating tolerance, while conflicting, suggests this may also hold true in the context of pain^80–83^.

### Translational relevance

Our studies on postmortem tissues support the exciting idea that the kappa opioid receptor system may also reduce pain transmission in humans. We show that *OPRK1* is expressed within the human superficial dorsal horn. Based on recent RNA-sequencing studies, human *OPRK1* spinal neurons appear to be a combination of excitatory and inhibitory neurons^50^, as we establish here in mice. In alignment with our finding that the majority of *OPRK1* neurons contain *TAC1* in human, this sequencing study also reported that *OPRK1* and *TAC1* are found in common excitatory spinal neuron clusters. Furthering the idea that spinal kappa opioid signaling may suppress pain in humans, the endogenous kappa opioid *PDYN* is also readily detected in the human spinal cord^27,50,76^. These data are important because they establish the kappa opioid receptor system (i.e., both the receptor and endogenous neuropeptide ligand) is present in the human spinal cord, and thus positioned to inhibit cell-types that encode pain. Future studies examining the effects of KOR signaling on spinal neuron function will be especially insightful in understanding the extent of translational relevance of our current findings.

Regardless of the mechanism of spinal KOR inhibition of pain, these results raise the question of whether spinal KOR may represent a promising avenue for the clinical treatment of pain. Centrally-acting kappa opioid receptor agonists have lost momentum in clinical development for pain out of concern for dysphoric and hallucinogenic side effects and instead peripherally-restricted agonists have garnered interest^8,84^. However, this central hurdle may be overcome through the use of biased KOR agonists such as nalfurafine that are well-tolerated by patients and lack abuse liability^78,79,85^.

### Conclusion

In summary, this study provides an in-depth characterization of the spinal neurons that express KOR as well as a model for how kappa opioid receptor signaling could block the spinal transmission of pain. Based on our suite of complementary rodent behavioral and molecular data, we propose KOR signaling suppresses pain by inhibiting the activity of excitatory neurons, and in turn reduces the release of pain-promoting neuropeptides such as *Tac1* within the spinal cord. Further, our data are translationally relevant and suggest an analogous mechanism may be at play within the human spinal cord. Taken into context with previous studies examining the kappa opioid system in the periphery and brain circuits, our work now allows for a holistic view of how KOR signaling modulates pain across the nervous system.

## Methods

### Animals

Animals were cared for in compliance with the National Institutes of Health guidelines, and experiments were approved by either the Medical College of Wisconsin (protocol number 00008654) or University of Pittsburgh (protocol number 21100045) Institutional Animal Care and Use Committees. Adult C57BL/6 mice were obtained from Charles River (strain 027) and allowed to acclimate to the vivariums for at least one week before beginning experimentation. *Oprk1^Cre^* mice (Jax, 035045) were generated and provided by Dr. Sarah Ross^32^ and maintained on a C57BL/6 background. Experiments were conducted on both male and female mice and experiments began when mice were 5-8 weeks of age. For all studies, littermates were randomly assigned to experimental groups. Mice were housed in a 12/12 hour light/dark cycle with food and water available ad libitum; nestlets and plastic housing domes were provided for enrichment.

### Donors

Human spinal cord tissue was obtained from the NIH NeuroBioBank at the University of Maryland, Baltimore in compliance with the Medical College of Wisconsin Institutional Review Board. Spinal cord tissue from the lumbar spinal segments (L3-L5) was collected from adult human organ donors (donor 1: white 43-year-old male, donor 2: 32-year-old female), flash frozen, embedded in optimal cutting temperature compound, and stored at −80°C until use.

### Intraspinal viral injections

Mice were anesthetized with a ketamine-xylazine-acepromazine cocktail (95 mg/kg-4.8 mg/kg-0.95 mg/kg, i.p.). To prepare animals for surgery, ophthalmic ointment (Dechra, 12920060) was applied to the eyes, the back was shaved, and the local antiseptic betadine (Fisher Scientific, 19-061617) was applied to the skin, followed by ethanol. Using a scalpel, a skin incision was made over the T12–L3 vertebrae. Small scissors were used to cut through fascia, and #5 forceps were used to separate muscle from the T12–T13 vertebrae, exposing the intervertebral space above the left L3/L4 and L4/L5 spinal segments^33–35^. Laminectomies were avoided to minimize bleeding and spinal cord trauma. Using a NanoInject III Programmable Nanoliter Injector (Drummond Scientific Company, 3-000-207) mounted on a stereotax (Kopf, Model 942), a glass capillary was carefully lowered down 300 µm from the surface of the dura, taking care to leave the posterior spinal artery intact. A total of 500 nL of virus was delivered at each spinal segment at a flow rate of 5 nL/s. The glass capillary was left in place for 5 min after each injection before being withdrawn. The skin incision was closed with 6–0 vicryl suture (Henry Schein, 6545672). After surgery, mice were injected subcutaneously with 0.3 mg/kg buprenorphine and 5 mg/kg ketoprofen and allowed to recover on a heating pad. Behavioral tests began four weeks after viral injection to allow for maximal and stable viral expression. At the conclusion of behavioral studies, intraspinal viral injections were confirmed with histology (described in Immunohistochemistry). One Oprk1::mCherry mouse was excluded from analyses because no mCherry was detected within the lumbar spinal cord.

### Viruses

To visualize the spatial distribution of *Oprk1^Cre^* neurons within the dorsal horn, an adeno-associated virus coding for Cre-dependent nuclear-localized red to green fluorescent protein, AAV8-EF1a-nuc-flox(mCherry)-EGFP (Addgene, 112677-AAV8, a gift from Brandon Harvey^36^), was injected intraspinally. For chemogenetic studies, an AAV encoding for either Cre-dependent hM3Dq-mCherry, AAV8-hSyn-DIO-hM3Dq-mCherry (Addgene, 44361-AAV8, a gift from Bryan Roth^37^), or Cre-dependent fluorescent reporter, AAV8-hSyn-DIO-mCherry (Addgene, 50459-AAV8, a gift from Bryan Roth), was injected intraspinally. Viruses were delivered into the spinal cord undiluted, with titers ranging from 2.2×10^12 to 2.5×10^13 GC/mL.

### Behavioral Studies

Each behavioral study was performed on at least two different cohorts of mice. Experimenters were blind to all experimental conditions throughout data acquisition and analysis. All behavior experiments were performed during the light cycle between 9 a.m. and 6 p.m in a designated, temperature-controlled room. On the day of testing, the mice were allotted at least 30 min to acclimate to a Plexiglas chamber. Dividers were placed in between chambers so animals could not see one another. When assessing non-reflexive behaviors (e.g., responses to algogens and pruitogens; chemogenetically-evoked spontaneous behaviors), responses were recorded via bottom-up video recordings. Mice were placed on an elevated glass platform and recorded with a Sony HD Camera (HDR-CX405). During these experiments, the experimenter left the behavior room once the last animal was injected; only mice that were successfully injected were included in data analysis.

### Pharmacological agents to target KOR

Nalfurafine (Cayman Chemicals, 36795) was dissolved in 4% DMSO in sterile saline immediately prior to each experiment. Aticaprant (Cayman Chemicals, 35464) was initially dissolved in equal parts ethanol and cremophor (Sigma Aldrich, C5135), and once the mixture was completely dissolved, it was further diluted in saline (working solution (in %): 10 ethanol: 10 cremophor: 80 saline). Each agent or its corresponding vehicle was delivered intrathecally 30 min prior to assessing capsaicin- or formalin-evoked nocifensive behaviors.

### Intrathecal injections

At least 1 day prior to behavioral testing, the backs of mice were shaved to allow for better visibility during intrathecal injections. On testing days, awake, behaving mice were pinned by the pelvic girdle, and a 25 µL Hamilton syringe with a 30 G needle attachment was inserted between the L5 and L6 vertebrae and successful placement was indicated by an involuntary lateral tail movement upon needle insertion. Nalfurafine (40 µg in 5 µL), aticaprant (10 µg in 5 µL), or their corresponding vehicles were injected at a rate of 1 µL/sec. The needle was held in place for an additional 5 s post-injection to ensure complete drug delivery. Following successful injection, mice were returned to their individual behavior chambers and recorded for 30 min during the drug onset of action.

### Intradermal injections

At least 1 day prior to behavioral testing, the calves of mice were shaved to allow for better visibility during intradermal injections. On testing days, awake, behaving mice were gently restrained with surgical towels and a 25 µL Hamilton syringe with a 30 G needle attachment was used to deliver capsaicin or chloroquine intradermally into the calf skin.

### Capsaicin evoked-nocifensive behavior

For pharmacological studies targeting KOR, mice received a 10 µL intraplantar injection of capsaicin (0.01% dissolved in 0.5% Tween-80 and 10% ethanol in sterile saline, Sigma Aldrich, M2028) into the right hindpaw. Mice were then returned to their individual behavior chambers and recorded for 15 min. The duration of nocifensive behaviors, including site-directed licking, biting, guarding, or flapping of the injected hindpaw, was quantified. In chemogenetic experiments, 90 min after CNO administration, 10 µL of 0.001% capsaicin was delivered intradermally into the calf ipsilateral to the intraspinal viral injection, mice were recorded for 15 min, and the duration of site-directed licking was quantified.

### Formalin test

For pharmacological studies targeting KOR, mice received an intraplantar injection of 10 µL formalin (2% in saline; Sigma Aldrich, 252549) into the right hindpaw. Nocifensive behaviors were quantified for 60 min post-injection and were analyzed in separate phases: phase I (0-10 min) and phase II (10-60 min). The same hindpaw nocifensive behavior guidelines as described above for capsaicin were used for quantification.

### Chemogenetic studies

Based on previous publications^33,38,39^, CNO hydrochloride (Tocris, 6329) was dissolved in sterile saline and mice were treated with 5 mg/kg CNO intraperitoneally (i.p.). Behavioral testing was completed within 0.5 - 2.5 h of CNO injection. Baseline nociceptive withdrawal thresholds (heat, cold, mechanical) were measured 1 day before CNO administration. Following CNO administration, some behavioral assays were paired (spontaneous behavior and Hargreaves, cold sensitivity and chloroquine-induced itch, mechanical sensitivity and capsaicin-induced pain) to minimize the number of CNO injections experienced by an individual mouse.

### CNO-evoked spontaneous behavior

Spontaneous nocifensive and itch-related behaviors directed toward the hindlimb ipsilateral to the intraspinal viral injection were quantified in the 30-60 min window after CNO administration. Spontaneous nocifensive behaviors observed included: flinching of the hindpaw (reported as number of flinches); spontaneous itch-related behaviors observed included: biting of the hindlimb/calf (reported in seconds), as established in previous studies^40,41^.

### Radiant heat sensitivity (Hargreaves test)

Mice were acclimated on a glass plate held at 30°C (IITC). Beginning 70 min following CNO administration, a radiant heat source (active intensity =15%; intermittent intensity=5%) was applied to the glass beneath the hindpaw and latency to paw withdrawal was recorded^42^. Three trials were conducted on each paw, with at least 5 min between testing the opposite paw and at least 10 min between testing the same paw. A 20 s cut-off latency was set to avoid tissue damage. Trials were averaged to determine the withdrawal latency of each paw.

### Noxious heat sensitivity (Hot Plate)

Beginning 40 min after CNO administration, each mouse was placed in a hot plate chamber (Columbus Instruments) held at 50°C, and the latency to response was recorded. “Response” was defined as either the first act directed towards the hindpaw, which was primarily licking or shaking, or an escape behavior such as jumping. The protocol was repeated at least 40 min later with the hot plate held at 54°C. A single trial was conducted at each temperature and a cut off latency of 60 s was used to prevent tissue damage.

### Cold sensitivity (Cold plantar assay)

Cold sensitivity was measured as previously described^43,44^. Mice were acclimated to a 1/4” glass plate, and a dry ice probe was made by packing finely crushed dry ice into a modified 3 mL syringe with a diameter of 1 cm. Beginning 30 min after CNO administration, the dry ice probe was applied to the glass beneath the plantar surface of the hindpaw and the latency to paw withdrawal was recorded. Three trials were conducted on each hindpaw, with 5 min between trials on opposite paws, and 10 min between trials on the same paw. A cut-off latency of 20 s was used to prevent tissue damage. Withdrawal latencies for each paw were determined by averaging values across trials.

### Mechanical sensitivity (von Frey test)

Mechanical sensitivity was measured using the simplified up-down (SUDO) method of the von Frey test^45^. Beginning 35 min following CNO administration, calibrated von Frey filaments (North Coast Medical Inc.) were applied to the plantar surface of the hindpaw for 2 s or until a paw withdrawal response occurred. For each trial, a series of five filament applications was performed to measure the paw withdrawal threshold. Three trials were conducted on each hindpaw, with 5 min between trials on opposite paws, and 10 min between trials on the same paw. Paw withdrawal thresholds for each paw were determined by averaging across trials and are reported in units of pressure (g/mm^2^) to account for differences in filament surface areas.

### Chloroquine-evoked itch behavior

Chloroquine (100 µg in 5 µL sterile saline, Sigma, C6628) was administered intradermally into the calf ipsilateral to intraspinal viral injection 75 min after the injection of CNO. Mice were video recorded for 30 min following chloroquine injection, and the duration of site-directed calf biting was quantified off-line.

### Capsaicin-induced Fos expression

All animals used capsaicin-induced *Fos* experiments were handled for 3 days leading up to experimentation to decrease non-specific *Fos* expression. Each mouse was scruffed twice for 2-3 s, then held in cupped palms until grooming behavior was observed. Mice were then wrapped in a small towel and gently restrained while an experimenter lightly tapped the plantar hindpaw with their index finger for 10 s to familiarize mice with the injection protocol. This was done a total of two times. On testing day, mice were injected with capsaicin or vehicle as described in “Capsaicin evoked-nocifensive behavior,” behavioral responses were recorded for 15 min, and spinal cords were harvested 30 min after injection and processed for RNAscope in situ hybridization.

### RNAscope fluorescence in situ hybridization (FISH)

Mice were anesthetized with isoflurane and rapidly decapitated. The L3-L5 spinal cord segments were harvested within 5 min, embedded in OCT, and flash frozen using 2-methylbutane chilled on dry ice. Tissue was kept on dry ice until cryosectioning. Mouse and human lumbar spinal cord samples were sectioned on a cryostat, and 15 μm sections were directly mounted onto Superfrost Plus slides (Fisher Scientific, 12-550-15). FISH experiments were performed according to the manufacturer’s instructions for fresh frozen samples (Advanced Cell Diagnostics, 323110), with minor species-specific modifications. Spinal cord sections were first fixed in ice-cold 4% paraformaldehyde for 15 min, then dehydrated in ethanol, permeabilized at room temperature with Protease IV for either 15 min (mouse) or 5 min (human), hybridized at 40°C with gene-specific probes to mouse or human (Table 1), and stored overnight in 5X saline sodium citrate. After rinsing tissues with wash buffer, the hybridized probe signal was amplified and fluorescently labeled using TSA Plus Fluorophores (Akoya Biosciences, NEL741001KT, NEL744001KT, and NEL745001KT). Slides were mounted with Prolong Gold with DAPI (Invitrogen, P36931) to visualize nuclei.

**Table 1.** Probes used for RNAscope analysis of mouse and human spinal cord. Mm, mus musculus; Hs, homo sapiens.

| **Target** | **Cat #** |
| --- | --- |
| *Mm-Oprk1* | 316111 |
| *Mm-Slc17a6* | 319171 |
| *Mm-Slc32a1* | 319191 |
| *Mm-Fos* | 316921 |
| *Mm-Tac1* | 410351 |
| *Mm-Npff* | 479901 |
| *Mm-Grp* | 317861 |
| *Mm-Tac2* | 446391 |
| *Mm-Cck* | 402271 |
| *Mm-Nts* | 420441 |
| *Hs-OPRK1* | 556711 |
| *Hs-TAC1* | 310711 |

### Immunohistochemistry

Mice were anesthetized with urethane and perfused with 4% paraformaldehyde. The lumbar spinal cord and dorsal root ganglia were dissected and postfixed in 4% paraformaldehyde at 4°C for 2 h. Tissues were washed in PBS, cryoprotected in 30% sucrose, then sectioned at 40 μm (spinal cord) or 20 μm (dorsal root ganglia) and directly mounted on Superfrost Plus slides. Tissues were incubated in a blocking solution consisting of 10% normal donkey serum (Jackson ImmunoResearch, 017-000-121) and 0.3% Triton X-100 solution (Sigma, 93443) in PBS for 1 h at room temperature. The primary antibodies rabbit anti-RFP (Rockland, 600-401-379, 1:1000) and rabbit anti-GFP (Invitrogen, A-11122, 1:1000) were diluted in antibody buffer consisting of 5% normal donkey serum and 0.3% Triton X-100 in PBS. Tissues were incubated in primary antibodies overnight at 4°C, then washed in PBS three times for 10 min. The secondary antibodies donkey anti-rabbit Alexa Fluor 555 (Invitrogen, A-31572, 1:500) and donkey anti-rabbit Alexa Fluor 488 (Invitrogen, A-21206, 1:500) were diluted in antibody buffer and applied for 1 h. Finally, tissues were washed again in PBS three times for 10 min and mounted with Prolong Gold with DAPI.

### Image acquisition and analysis

Sections were imaged at full tissue thickness using an upright epifluorescent microscope (Nikon Eclipse 80i with an Nikon Plan Apo 10X air objective), or a slide scanner (Olympus VS200X, 20X air objective). All analysis was completed off-line using FIJI (ImageJ, NIH) or QuPath software. For FISH experiment quantification, three spinal cord hemisections were manually quantified from each mouse, and two spinal cord hemisections were quantified from each human donor. A positive cell was defined as a cell with a clearly defined nucleus and at least 5 fluorescent puncta around the nucleus. To confirm viral expression in *Oprk1^Cre^* mice, every 4th section was evaluated for mCherry or EGFP expression. On average, unilateral intraspinal viral injections resulted in 3 mm of rostral-caudal viral spread spanning the L3-L5 segments.

Analyses of the spatial distribution of spinal neurons in mouse and human used the following boundaries: In mouse, the superficial dorsal horn (SDH) was visually defined as the region from the surface of the dorsal horn gray matter to the bottom of the substantia gelatinosa, roughly corresponding to laminae I and II_o_. In human, the SDH was defined as the region bounded by the surface of the dorsal horn gray matter and the bottom of the lamina II, as indicated by a dense banding of nuclei between lamina II and III. In both mouse and human, the deep dorsal horn (DDH) consisted of the remaining dorsal horn lamina, roughly III–VI. Lateral spinal nucleus (LSN) neurons were defined as those located lateral to the dorsal horn gray matter, within the dorsolateral funiculus. For human tissue samples, a gamma correction was applied to enhanced visibility of DAPI.

### Statistical analysis and data visualization

Microsoft Excel, R, and GraphPad Prism were used for data organization, visualization, and statistical analyses. Statistical analyses were performed on a per mouse basis. Prior to running analyses, we first assessed whether data were normally distributed. For datasets comparing two independent groups with n ≥ 8 mice, we performed a D’Agostino-Pearson test; otherwise, a Shapiro-Wilks test was performed. If each group was normally distributed, data were analyzed with an unpaired Student’s t-test; if one or more groups were not normally distributed, data were analyzed with a Mann-Whitney test. For datasets assessing the relationship between two factors, normality was assessed using QQ plots, and data were analyzed with a 2-way ANOVA. Across statistical tests, a Holm-Sidak test was used to correct for multiple comparisons when appropriate. Throughout the figures, statistical significance is indicated with the following symbols: *P < 0.05, **P < 0.01, ***P < 0.001, and ****P < 0.0001. Experiment--specific details can be found in the figure legends. Data are presented as mean ± SEM, with open circles representing individual mice or humans.

## Author contributions

Conceptualization: T.D.S.

Data curation: N.A.K, T.D.S.

Formal analysis: N.A.K, B.M.W., T.D.S.

Funding acquisition: T.D.S.

Investigation: N.A.K, B.M.W., L.G.W., L.G.F., A.P.M., T.D.S.

Projection administration: T.D.S.

Supervision: T.D.S.

Visualization: N.A.K, B.M.W., L.G.W., T.D.S.

Writing - original draft: N.A.K, B.M.W., L.G.W., T.D.S.

Writing - review & editing: N.A.K, B.M.W., L.G.W., T.D.S.

## Acknowledgements

We thank Drs. Cheryl Stucky, Harrison Stratton, and Jonathan Enders for their thoughtful comments and suggestions on our manuscript, Dr. John McCorvy for providing nalfurafine, and Dr. David Baranger for statistical consultation. We thank the Medical College of Wisconsin (MCW) Cardiovascular Research Center Microscopy Core (RRID:SCR_028523) for access to microscopes used for RNAscope tissue imaging and the Biomedical Resource Center staff at MCW for providing animal husbandry and veterinary care. We also thank the NIH NeuroBioBank, as well as the organ donors and their loved ones, for providing access to human tissue samples. Select figure panels were created using icons from Biorender. This work was funded in part by National Institutes of Health Grant R00 NS126569 to T.D.S., startup funds to T.D.S. from MCW, as well as the MCW Engineering Core (RRID:SCR_028303, grant S10OD032136 to the Medical College of Wisconsin), which fabricated behavioral equipment.

## Declaration of generative AI and AI-assisted technologies in the writing process

No generative AI or AI-assisted technologies were used in the preparation of this work.

