## Supplemental Data for "Kappa opioid receptors in excitatory spinal neurons gate acute pain: evidence from mouse and human"

Nicole A. Kooij *et al*.

**This file includes:**

Figures S1 to S3

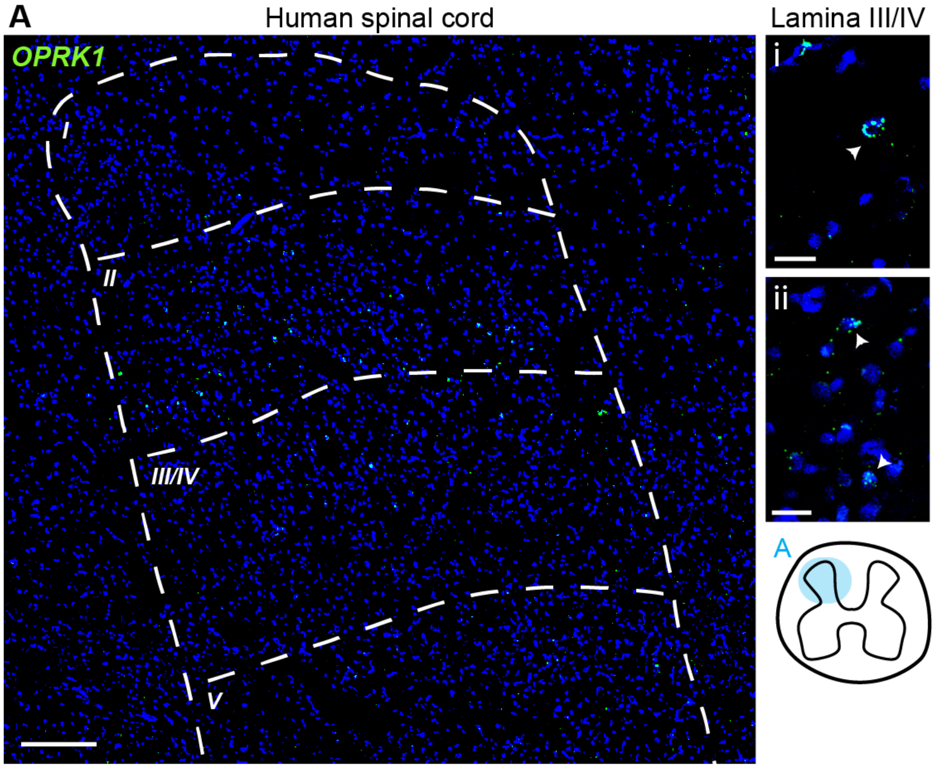


**Supplemental Fig. 1 (related to Fig. 2). Distribution of *OPRK1*-expressing neurons in the deep dorsal horn of human spinal cord**

(**A**) Representative FISH image of *OPRK1*$^{+}$ neurons distributed throughout the human dorsal horn, with insets highlighting *OPRK1* expression within deep dorsal horn neurons. Scale bars: A, 250 μm; $A_i$ and $A_{ii}$, 50 μm.

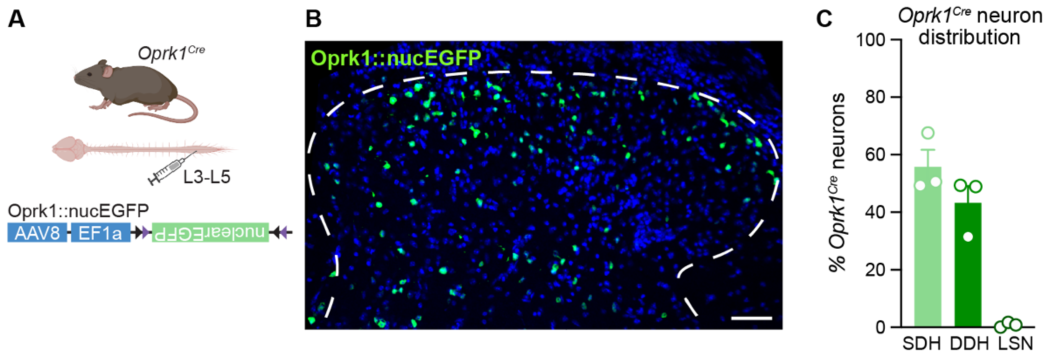


**Supplemental Fig. 2 (related to Fig. 3). Distribution of *Oprk1*$^{Cre}$-expressing neurons in the mouse spinal cord dorsal horn.**

(**A**) Strategy to selectively visualize *Oprk1*$^{Cre}$ spinal neuron cell bodies. *Oprk1*$^{Cre}$ spinal neurons have extensive processes (see Fig. 3B). Thus, to more readily visualize discrete *Oprk1*$^{Cre}$ neurons within the dorsal horn, the Cre-dependent virus AAV8-EF1a-Nuc-flox(mCherry)-EGFP (Oprk1::nucEGFP) was injected unilaterally into the lumbar spinal cord of *Oprk1*$^{Cre}$ mice to drive localized expression of EGFP within the nucleus.

(**B**) Representative image and

(**C**) quantification of the percentage of *Oprk1*$^{Cre}$-expressing neurons distributed throughout the SDH, DDH, and LSN. Scale bar, 50 μm (N=3 mice; n=180-288 total *Oprk1*$^{Cre}$ neurons per mouse). Data are shown as mean ± SEM, with open circles representing individual mice.

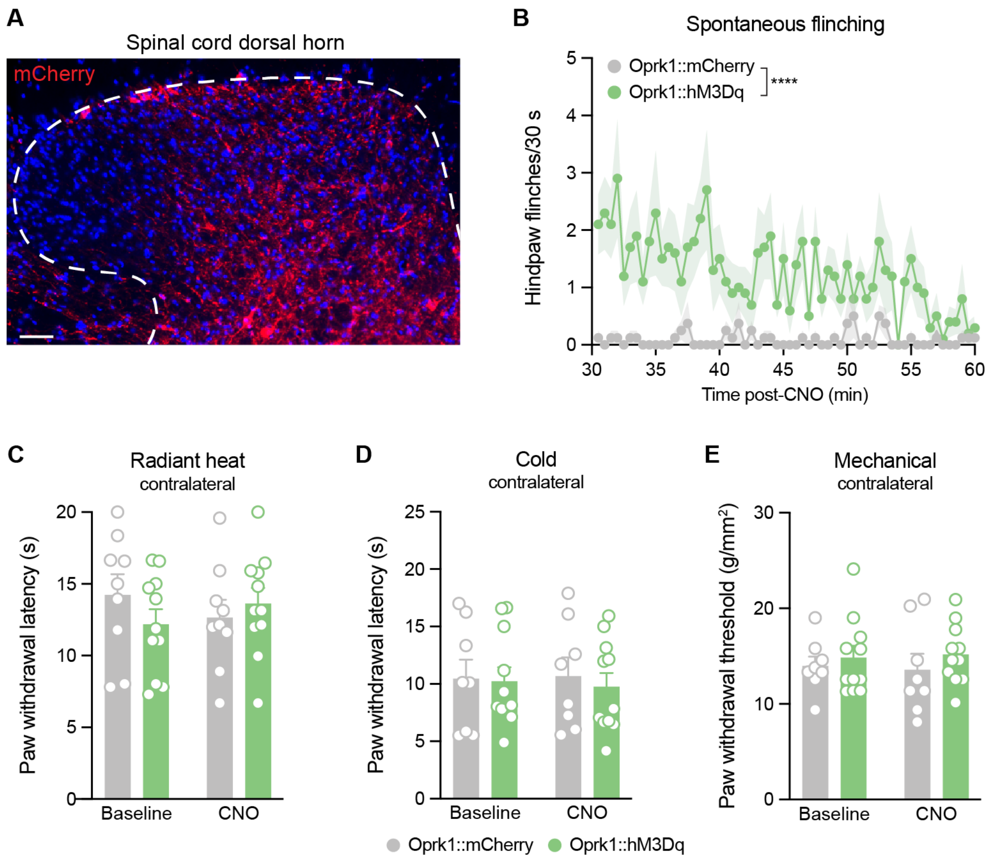


**Supplemental Fig. 3 (related to Fig. 3). Extended analysis of nocifensive behaviors and nociceptive withdrawal thresholds following *Oprk1^Cre* spinal neuron activation.**

(**A**) Additional representative image of virally-mediated recombination in the spinal cord of *Oprk1^Cre* mice, as indicated by mCherry expression. Image representative of N=3 mice. Scale bar, 50 µm

(**B**) Oprk1::hM3Dq mice display a greater number of hindpaw flinches following systemic CNO administration (5 mg/kg, i.p.) as compared to Oprk1::mCherry mice, binned over time in 30 s windows (n = 8-10 mice per group). 2-way RM ANOVA, main effect of virus: ****$p<0.0001$. Cumulative data are presented in Fig. 3D. Of note, while we observed a significant increase in the number of spontaneous flinches, flinches were intermittently spaced (average of < 3 flinches/30 s) and peaked between 30-40 min after CNO administration. Thus, the spontaneous flinches were unlikely to have interfered with measures of nociceptive withdrawal thresholds.

(**C**) Chemogenetic activation of *Oprk1^Cre* spinal neurons did not affect the contralateral hindpaw withdrawal thresholds of Oprk1::hM3Dq mice to radiant heat (n = 9-11 mice per group),

(**D**) cold (n = 8-11 mice per group), or

(**E**) mechanical (n = 8-11 mice per group) stimuli relative to Oprk1::mCherry controls.

B-D: Data were analyzed with a 2-way RM ANOVA. Data are shown as mean ± SEM, with closed circles representing group means and open circles representing individual mice.
